# SDC4 drives fibrotic remodeling of the intervertebral disc under altered spinal loading

**DOI:** 10.1101/2025.03.13.643128

**Authors:** Kimheak Sao, Makarand V. Risbud

## Abstract

Alterations in physiological loading of the spine are deleterious to intervertebral disc health. The caudal spine region Ca3-6 that experiences increased flexion, showed disc degeneration in young adult mice. Given the role of Syndecan 4 (SDC4), a cell surface heparan sulfate proteoglycan in disc matrix catabolism and mechanosensing, we investigated if deletion could mitigate this loading-dependent phenotype. Notably, at spinal levels Ca3-6, *Sdc4-*KO mice did not exhibit increased collagen fibril and fibronectin deposition in the NP compartment or showed the alterations in collagen crosslinks observed in wild-type mice. Similarly, unlike wild-type mice, NP cells in *Sdc4*-KO mice retained transgelin (TGLN) expression and showed absence of COL X deposition, pointing to the preservation of their notochordal characteristics. Proteomic analysis revealed that NP tissues responded to the abnormal loading by increasing the abundance of proteins associated with extracellular matrix remodeling, chondrocyte development, and contractility. Similarly, downregulated proteins suggested decreased vesicle transport, autophagy-related pathway, and RNA quality control regulation. Notably, NP proteome from *Sdc4* KO suggested that increased dynamin-mediated endocytosis, autophagy-related pathway, and RNA and DNA quality control may underscore the protection from increased flexion-induced degeneration. Our study highlights the important role of SDC4 in fine-tuning cellular homeostasis and extracellular matrix production in disc environment subjected to altered loading.

## INTRODUCTION

Intervertebral disc degeneration is a major contributor to chronic low back pain and is characterized by significant alterations to the disc extracellular matrix (ECM)(1, 2). The disc is composed of three distinct tissue compartments: the central nucleus pulposus (NP), the annulus fibrosus (AF) that surrounds the NP, and the endplates (EP) that cap the disc on the superior and inferior side. The NP is rich in proteoglycans, primarily aggrecan (ACAN), essential for maintaining tissue hydration, resilience, and compressive strength. The AF consists of concentric layers of collagen I and II fibers, which provide tensile strength and structural stability by effectively counteracting tensile stress. Whereas, the EPs form the critical barrier and interfaces with adjacent vertebrae to facilitate nutrient exchange and contribute to overall biomechanical integrity(3). During degeneration, whether due to genetic predisposition, aging, or aberrant mechanical stress, there is a progressive loss of water-attracting proteoglycans with a concomitant increase in collagen and fibronectin (FN) deposition. Accordingly, altering ECM composition diminishes the ability of the disc to absorb and distribute loads, resulting in altered cellular behavior and compromised mechanical performance. Thus, preserving the integrity of ECM is crucial for maintaining disc health(4).

A recent study showed that in the mouse spine, the caudal region between Ca3-6 experiences accelerated degenerative changes due to increased flexion associated with posture(5). These discs in young adult mice show early onset fibrotic changes, making them an important natural model to investigate the mechanisms underlying loading-induced disc degeneration. Syndecan 4 (SDC4), a cell surface, transmembrane heparan sulfate proteoglycan, is well-known for its role in cell signaling, ECM remodeling, and inflammatory responses(6–9). SDC4 is enriched in focal adhesions and interacts with cell surface integrins, the primary transmembrane mechanosensing receptors, to modulate their function(10–12). A recent study showed that SDC4 modulates cell mechanics in response to localized tension by activation of EGFR and β1-integrin(13). We have earlier shown an elevated SDC4 expression in human degenerated disc tissues. Moreover, pro-inflammatory cytokines TNF-α and IL-1β regulate SDC4 expression, which plays a key role in the pathogenesis of disc degeneration by promoting aggrecan degradation via ADAMTS-5 dependent mechanism(9). Despite the known contribution of SDC4 in cell mechanosensing in other cell types, its role in loading-induced disc degeneration is not known.

In this study, we investigated whether loss of SDC4 can mitigate the early-onset disc degeneration in mice evident in the caudal spine at levels Ca3/4–5/6, which naturally experience increased flexion. We found that SDC4 loss impacted aberrant ECM remodeling, preserved notochordal characteristics of NP cells, and affected broader processes linked to cytoskeletal rearrangement, ciliary function, dynamin-mediated endocytosis, and autophagy. We demonstrate that SDC4 is an essential constituent of the molecular machinery that transduces and links the altered mechanical loading on intervertebral disc to cellular responses.

## RESULTS

### SDC4 mediates early-onset disc degeneration in the mouse proximal caudal spine that naturally experiences increased flexion

To investigate whether SDC4 contributes to disc tissue remodeling in response to altered flexion and sustained loading changes, we chose to investigate caudal levels Ca3/4, Ca4/5, and Ca5/6 in young adult mice as a model of mechanical load-induced disc degeneration(5). We hypothesized that since SDC4 interacts with integrins and contributes to aspects of cellular mechanosensing and matrix homeostasis, deletion will delay or mitigate early signs of degeneration (Fig. 1A). We performed a visual assessment of micro-dissected tissues from discs Ca3/4, Ca4/5, and Ca5/6 from 6-month-old wild-type (WT) and *Sdc4* knockout (KO) mice and compared them to discs from a contiguous spine region Ca6-9 that does not experience altered loading. Visual inspection of NP tissue consistency in wild-type mice revealed that gelatinous NP tissue in regions with increased flexion was replaced by fibrous tissue. On the contrary, the gelatinous appearance of NP tissue was preserved in *Sdc4*-KO mice (Fig. 1B, B’). A level-by-level analysis showed altered consistency in 33% of Ca3/4, 83% of Ca4/5, and 78% of Ca5/6 discs in WT mice and 19%, 38% and 38% corresponding discs in KO mice, representing a significant degeneration at each of the level compared to KO mice (Fig. 1B”). We then performed a histological assessment of disc morphology using the modified Thompson grading scheme. Safranin O/Fast Green/Hematoxylin staining of the Ca3-6 disc sections confirmed significant signs of degeneration in WT mice. Particularly, degeneration was characterized by a loss of vacuolated NP cells, replacement of proteoglycan-rich matrix with fibrotic tissue, NP tissue clefts, and AF lamellar buckling. In contrast, a higher proportion of KO discs showed healthy cell and matrix morphology without any noticeable degenerative structural changes (Fig. 1C). On the other hand, irrespective of the genotype, Ca6-9 discs had a healthy morphology (Fig. 1C). Accordingly, NP and AF compartment degeneration scores for Ca3-6 discs were significantly higher in wild-type compared to KO mice (Fig. 1C’). Notably, more than 50% of WT NP showed severe degeneration and were consistently scored as grade 4, in contrast to KO NP tissue with a grade score of 1 (Fig. 1C”). However, Ca6-9 discs showed low and comparable Thompson scores between the WT and KO mice. These findings suggested that spinal levels Ca3/4–5/6 are prone to early disc degeneration as a consequence of altered mechanical environment resulting from increased spine flexion.

**Fig. 1.**
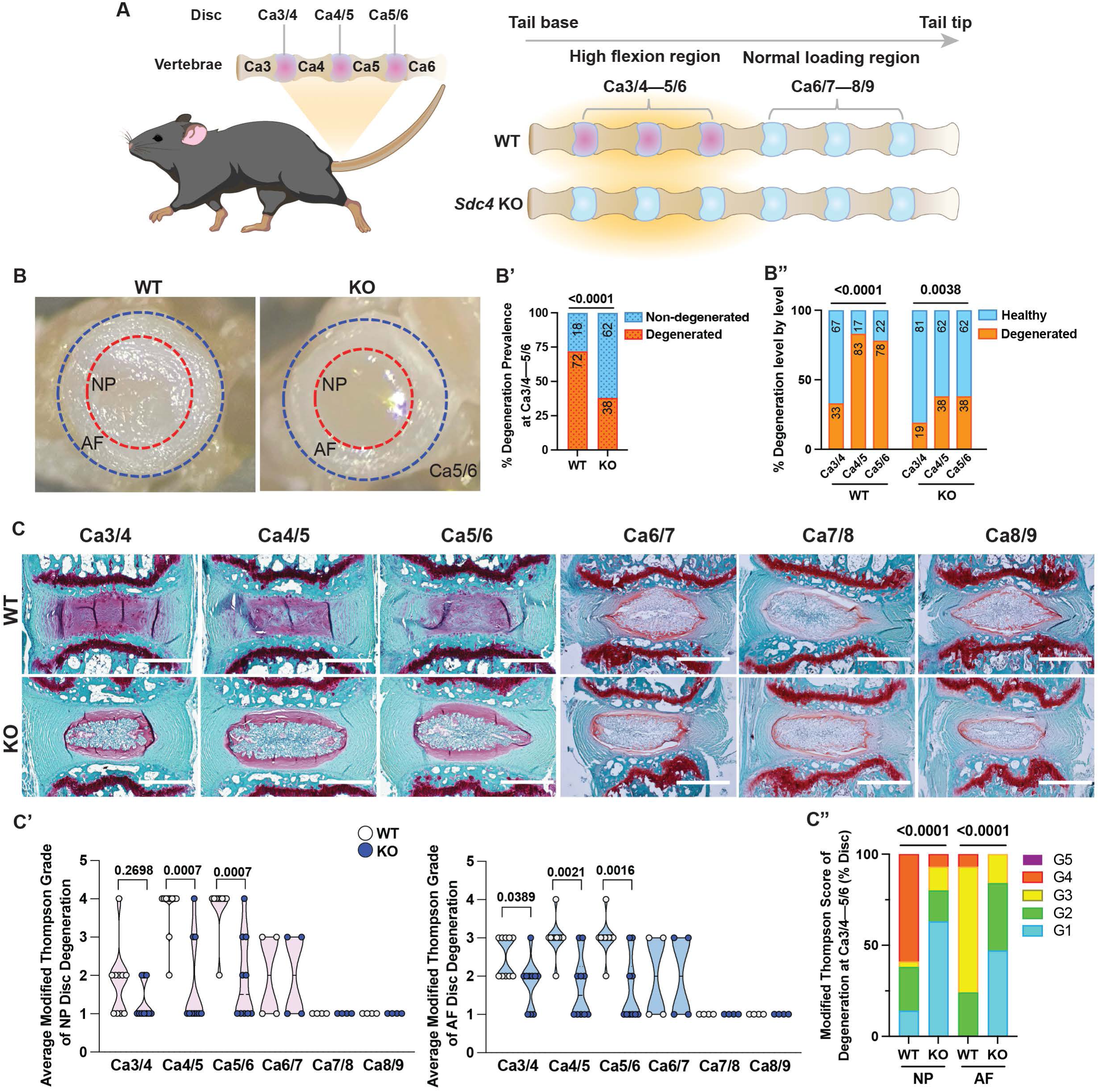
SDC4 mediates early onset of degeneration in discs experiencing increased flexion. (A) Cartoon illustration of the mouse caudal spine depicting proximal discs Ca3/4–5/6 that experience high flexion and evidence early degeneration (pink discs). Discs between Ca6-9 experience normal physiological loading (blue discs). We hypothesized that loss of SDC4 will delay this early degeneration phenotype in SDC4 KO mice (discs shown in blue). (B) Representative pictures showing gross appearance of the NP and AF tissues of caudal discs experiencing high flexion. (B’) Incidence of degeneration in WT and KO mice. (B”) Proportions of Ca3/4–5/6 discs with degeneration in WT and KO mice. 18 WT mice (10M, 8F), 54 discs; 21 KO mice (10M, 11F), 63 discs. Significance was determined using a χ^2^ test. (C) Representative Safranin-O/Fast Green/Hematoxylin staining showing morphology of Ca3/4–8/9 discs. (C’) Modified Thompson Grading (Grade 1-5) of caudal discs from WT and KO mice. (C”) Distribution of Modified Thompson scores. For Ca3-6, 10 WT mice (7M, 3F), 29 discs; 10 KO mice (9M, 1F), 30 discs. For Ca6-9, 4M WT and KO mice, 12 discs/genotype. Scale bar, 250μm. Significance was determined using either unpaired t-test or χ^2^ test where graphs represent contingency plots. Violin plots show score distribution with median and quartile range, P < 0.05.

### SDC4 promotes collagen crosslinking in disc in response to increased spinal flexion

To examine collagen fiber structure and thickness changes accompanying early onset disc degeneration due to increased spinal flexion, we used picrosirius red staining with polarized microscopy (Fig. 2A). While birefringence analysis of AF collagen fibrils showed no differences between genotypes (Fig. 2A’), approximately 59% of WT discs, compared to 20% discs in *Sdc4*- KO mice, showed abnormal collagen deposition in the NP compartment (Fig. 2A”). At this age, discs of both WT and KO mice from a contiguous spine region that experience normal loading do not show collagen deposition in NP (Suppl. Fig.1). NP collagen fibril thickness of discs that exhibited fibrosis was assessed and showed no genotype-specific differences (Fig. 2A”). Next, we used a collagen hybridization peptide (CHP) binding assay to ascertain the extent of collagen denaturation and observed a significantly higher abundance of denatured collagen in both NP and AF of WT mice. In contrast, *Sdc4*-KO mice showed very little binding of CHP, suggesting a lack of collagen denaturation (Fig. 2B and B’). Additionally, unlike WT mice, we noted a substantially lower levels of fibronectin (FN) in *Sdc4*-KO discs (Fig. 2C).

**Fig. 2.**
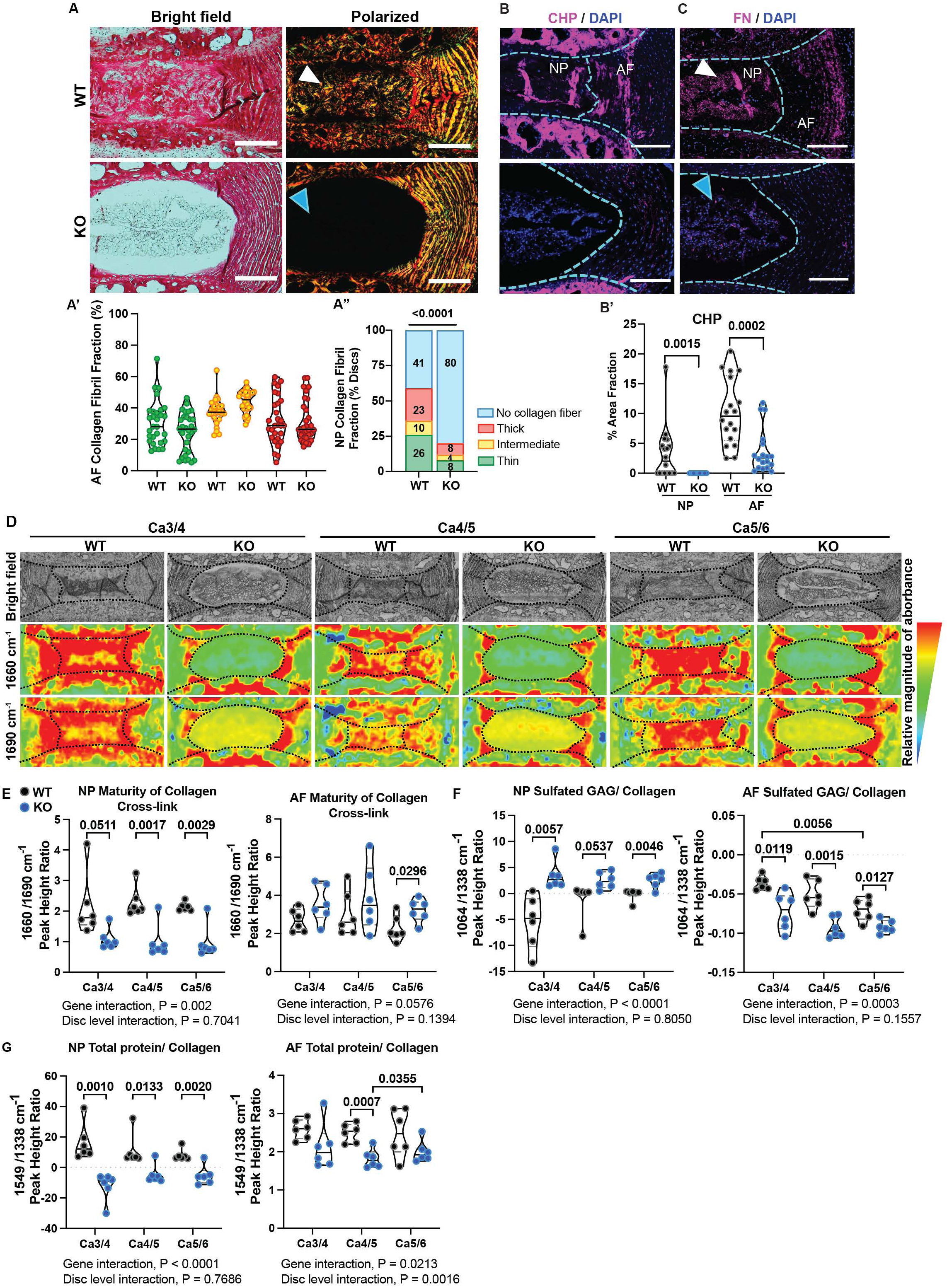
SDC4 increases collagen crosslinking in discs experiencing increased flexion. (A) Brightfield and polarized images of Picrosirius Red stained caudal discs. (A’) Quantification of AF collagen fibril fraction, showing thin (green), medium (yellow) and thick (red) fibers. (A”) Incidence of NP fibrosis and collagen fibril composition. For Ca3-6, 10 WT mice (7M, 3F), 29 discs; 10 KO mice (9M, 1F), 30 discs total. Scale bar, 250μm. (B) Representative images of collagen hybridization peptide (CHP) binding in the WT and KO disc. Scale bar, 250μm. Quantification of CHP signal in WT and KO Ca3-6 discs; 6 mice/genotype (5M, 1F), 18 discs. Scale bar, 250μm. (C) Representative immunostaining images of FN deposition in the disc, scale bar, 200μm. (D) Representative brightfield images of discs with corresponding chemical spectral maps at 1660 and 1690cm^-^¹. The color scale indicates the relative absorbance magnitude. (E-G) Peak height ratios showing chemical signatures characterizing the ECM. 6M WT and KO mice, 18 discs/ genotype. Significance was determined using Two-way ANOVA with Tukey’s multiple comparison test (E-G). Significance was determined using either Welch’s t-test or Mann-Whitney test (A’, B’) or χ^2^ test (A”). Violin plots show score distribution with median and quartile range, P < 0.05

To further assess collagen cross-linking, sulfated glycosaminoglycan (sGAG) composition and collagen content, imaging-Fourier Transform Infrared Spectroscopy (FTIR) was used to analyze the chemical composition of the discs. The Ca3-6 disc sections were imaged, and a second derivative function was applied to enhance the resolution of overlapping peaks.

Key peaks analyzed centered around 1660cm^-1^ (amide I, pyridinoline [PYR]), 1690cm^-1^ (dehydro-dihydroxynorleucine [de-DHLNL]), 1064cm^-1^ (chondroitin sulfated GAG specific peak), 1338cm^-1^ (collagen specific peak), and 1549cm^-1^ (amide II). The peak height intensity of mature non-reducible trivalent crosslinks (PYR) at 1660cm^-1^ was compared to that of immature divalent precursor crosslinks (de-DHLNL) at 1690cm^-1^ in the NP and AF to determine relative abundance of maturity of collagen crosslinks. Peak height ratio revealed that NP tissues from wild-type mice possessed significantly higher collagen crosslink maturity compared to *Sdc4* KO (Fig. 2D-E).

Interestingly, maturity of AF collagen crosslinks in *Sdc4*-KO and WT was similar except at Ca5/6, with a higher mature collagen cross-linking in KO mice (Fig. 2E). We speculated that this higher cross-linking at Ca5/6 in KO mice may be due to decreased collagen turnover. Next, spectral peak height ratios at 1064/1338cm^-1^ and 1549/1338cm^-1^ were used to determine the relative abundance of sGAGs and total protein/collagen content, respectively. Our results showed higher relative sGAG in NP tissues of *Sdc4*-KO across all levels, whereas WT AF possessed higher sGAG than KO (Fig. 2F). Furthermore, using 1549/1338cm^-1^, we found that WT NP had more relative total protein/collagen, while KO AF had lower baseline total protein/collagen (Fig. 2G). Overall, our results suggested that in response to increased spinal flexion, SDC4 mediates fibrotic remodeling of NP tissue by increasing collagen cross-linking and FN deposition.

### SDC4 promotes disc ECM production and fibrotic remodeling in response to increased spinal flexion

Given that SDC4 loss results in the preservation of ECM architecture and composition in the discs experiencing altered loading, we studied localization and abundance of select disc ECM molecules by immunohistochemistry. We noted an increased abundance of collagen I (COL I) in AF compartment of WT compared to KO discs, with no discernable difference in the NP (Fig. 3A-A’). Notably, collagen II (COL II) staining revealed higher abundance in the NP of WT than *Sdc4*-KO mice (Fig. 3B-B’), suggesting that the observed fibrosis primarily comprises of thin COL II fibrils. Moreover, dysregulation of the mechanical environment caused NP cells to adopt a hypertrophic phenotype evidenced by the increased collagen X (COL X) deposition only in wild-type mice. Similarly, NP cells in wild-type mice showed a loss of progenitor-like characters noted by decreased transgelin (TAGLN) abundance (Fig. 3C-D’). To determine whether the hypertrophic differentiation of NP cells impacts ECM composition, we measured the key NP ECM constituents, aggrecan (ACAN) and chondroitin sulfate (CS). While the ACAN abundance as a percentage of NP area appeared higher in wild-type discs, it was likely a reflection of the reduction of the compartment size causing compaction of the matrix (Fig. 3E-E’). CS levels showed no significant differences between the genotypes (Fig. 3F-F’), but trended higher in the KO, a finding corroborated by FTIR analysis. Overall, our results showed that altered mechanical loading causes SDC4-dependent shift in cellular phenotype, transitioning from TAGLN-positive notochordal NP cells to the acquisition of a COL X-positive hypertrophic phenotype, promoting collagen-rich matrix deposition.

**Fig. 3.**
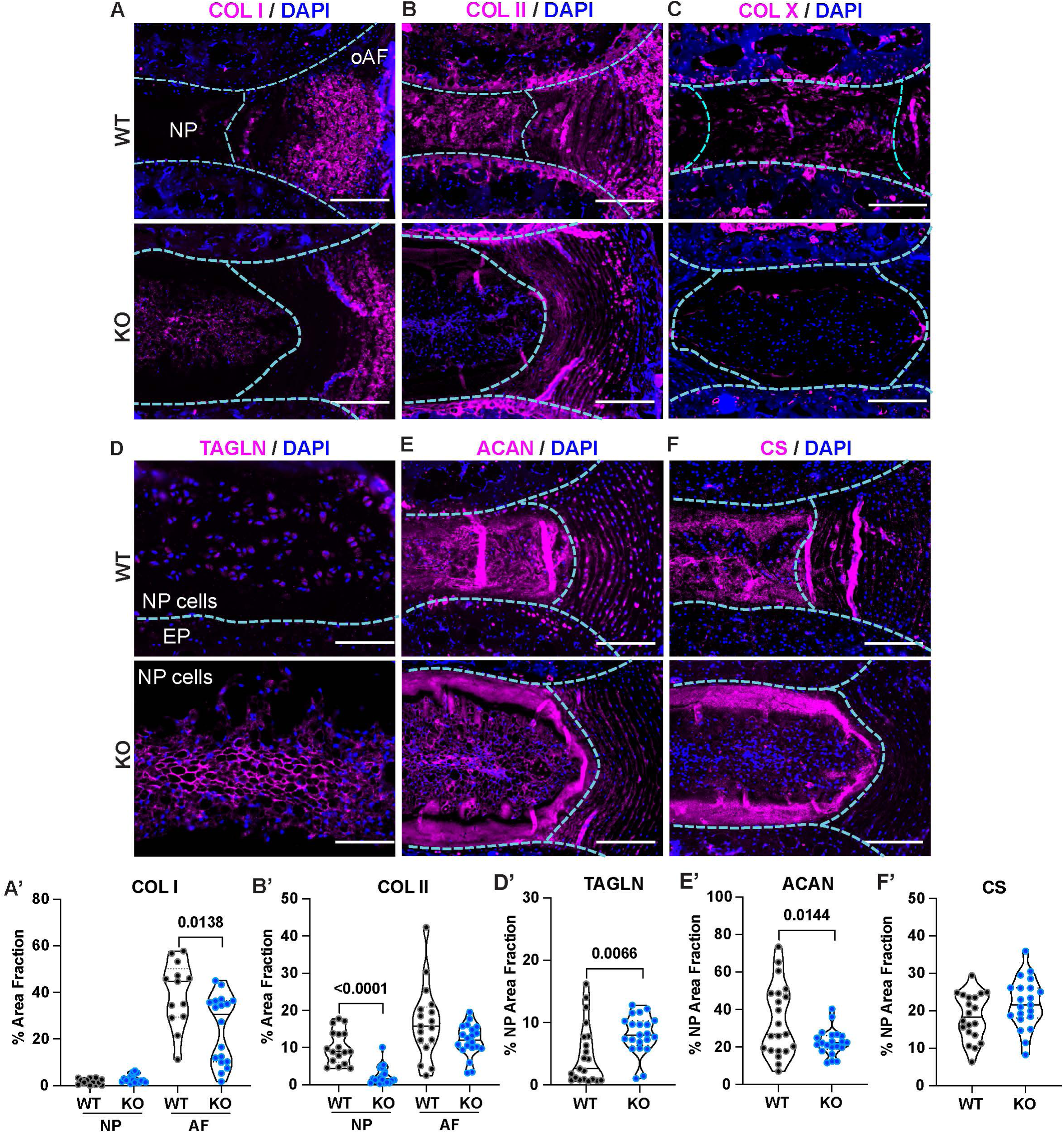
SDC4 contributes to ECM production and mediates fibrotic remodeling under condition of increased flexion-induced loading. (A) Representative immunostaining and quantification of (A-A’) COL I, (B-B’) COL II, (C) COLX, (D-D’) TAGLN, scale bars at 100µm, (E-E’) ACAN, (F-F’) CS. All other scale bars, 200µm. Per marker, both sexes were used. 4-7 WT mice, 13-21discs; 6-7 KO mice, 18-21 discs total. Dotted lines are used to demarcate NP, AF, and cartilaginous endplate (CEP) compartments. Violin plots show score distribution with median and quartile range. Significance was determined using an unpaired Welch’s t-test or Mann-Whitney test, P < 0.05.

### Increased spinal flexion accelerates matrix remodeling of the NP compartment with concomitant downregulation of cellular stress response

To gain broader mechanistic insights into how increased spinal flexion affects NP tissue health, we used label-free LC-MS/MS to obtain proteomic profiles of Ca3/4–5/6 discs and adjoining Ca6/7–8/9 discs; comparisons were performed within the same and across the genotypes (Fig. 4A, Suppl. Fig. 2A-A’). Using the data-independent acquisition neural network (DIA-NN) filtering method, a total of 4655 unique proteins were identified. To ensure consistency with a previously reported study(14), matrisome analysis was performed using Matrisome AnalyzeR(15). This method identified an average of 3822 proteins per sample, including both matrisome and non-matrisome proteins (min: 1768; max: 4500; median: 4155). Of these, an average of 225 proteins per sample were associated with matrisome (Fig. 4A’, Suppl. Fig. 2B, Table 2). Overall, our protein extraction and proteomic results align with the findings by Kudelko *et al.*(14).

**Fig. 4.**
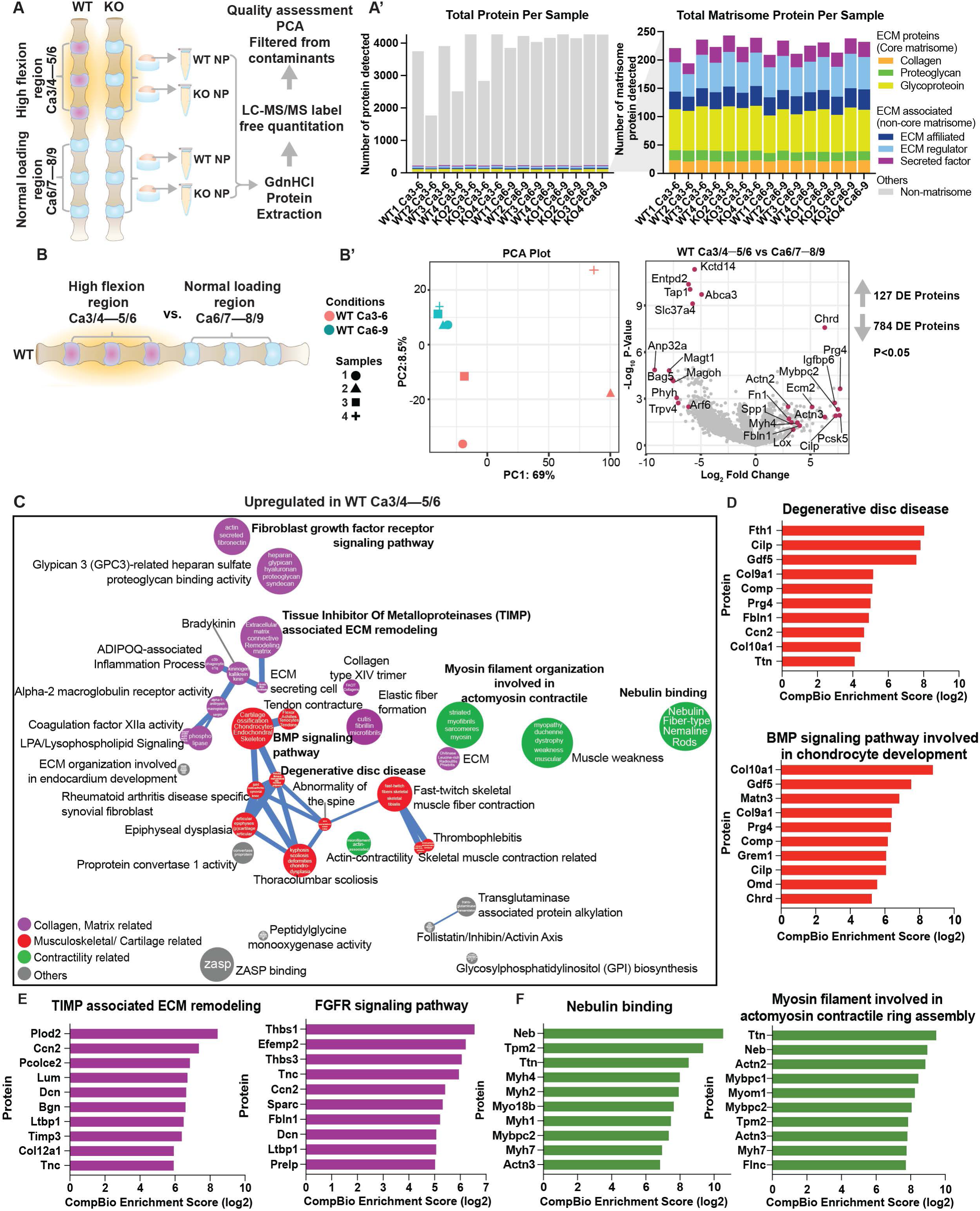
Proteomic analysis reveals NP ECM remodeling under increased flexion-induced loading in wildtype mice. (A) Cartoon illustration depicting the isolation and protein extraction process from NP tissues under increased flexion (Ca3/4–5/6) and normal loading (Ca6/7–8/9). (A’) Total proteins and matrisome-associated proteins detected from each sample. (B) Comparison of WT caudal discs under high flexion versus discs under normal loading condition. (B’) PCA plot shows sample cluster separation, and log^2^ fold change volcano plot highlights upregulated and downregulated DEPs. (C) Themes associated with upregulated DEPs under increase flexion-induced stress are highlighted. The size of a sphere is related to its enrichment score and the thickness of the lines connecting themes signifies the number of proteins shared between them. (D-F) CompBio analysis of DEPs and themes whose abundances were significantly upregulated under altered overloading.

To determine changes in proteome associated with increased flexion, we compared profiles of Ca3/4–5/6 to Ca6/7–8/9 in wild-type mice (Fig. 4B). Principle component analysis (PCA) revealed distinct clustering between two groups of samples. Differentially expressed protein (DEP) analysis identified 127 upregulated and 784 downregulated proteins (p ≤ 0.05, FC ≥1.5) (Fig. 4B’). We used the CompBio tool to analyze DEPs, identifying contextually relevant concepts and pathways, which were organized into themes (colored spheres) and clusters of related themes (colored spheres and connecting sticks). (Table 1).

From the CompBio analysis, we identified three prominent thematic superclusters that were upregulated in response to increased flexion: 1) collagen and matrix-related, 2) contractility-related, and 3) musculoskeletal/cartilage-related. The top DEPs with high CompBio normalized enrichment scores (NES) contributing to these superclusters were highlighted (Fig. 4C-F). Specifically, high flexion elevated levels of FTH1, CILP, GDF5, GREM1, COL9A1, COL10A1, COMP, and MATN3, which are associated with degenerative disc disease and bone morphogenetic protein (BMP) signaling involved in chondrocyte development themes (Fig. 4D).

Upregulation of PLOD2, CCN2, PCOLCE2, TIMP3, EFEMP2, THBS1/3, and LTBP1 in tissue inhibitor of matrix metalloproteinases (TIMP)-associated ECM remodeling and fibroblast growth factor receptor (FGFR) signaling pathway themes contribute to ECM organization, collagen cross-linking, and structural stabilization (Fig. 4E)(16–20). Contractility-related themes including nebulin binding and myosin filament involved in actomyosin contractility featured proteins such as NEB, TPM2, TTN, ACTN2, MYBPC1/2, and MYH1/2/4 (Fig. 4F)(21, 22).

NP tissue under high flexion exhibited downregulation of biological themes relating to vesicle transport, endoplasmic reticulum-associated protein degradation (ERAD), and mRNA quality control (Fig. 5A). DEPs associated with vesicle trafficking included CCZ1, PIKFYVE, VAC14, BECN1, FYCO1, GBF1, ARF6, CHMP2B, VPS4B, RHOG, RAB35, and CDC42 (Fig. 5B-C). Furthermore, the downregulation of EXOSC10, DIS3, PARN, PAN2, PIKFYVE, INPPL1, PNKP, and RECQL indicated impaired stress response pathways, including the RNA exosome complex (responsible for 3’ to 5’ exonucleolytic mRNA degradation), PI3K-mediated conversion of PIP2 to PIP3 (critical for cellular signaling), and DnaB helicase complex which is involved in DNA replication (Fig. 5D-E). These results underscored characteristics of cartilaginous tissue degeneration, where excessive matrix remodeling, coupled with imbalanced anabolic and catabolic activity, and overall impairment of the cellular ability to manage stress and maintain proper protein and RNA integrity contributed to structural deterioration and loss of tissue integrity.

**Fig. 5.**
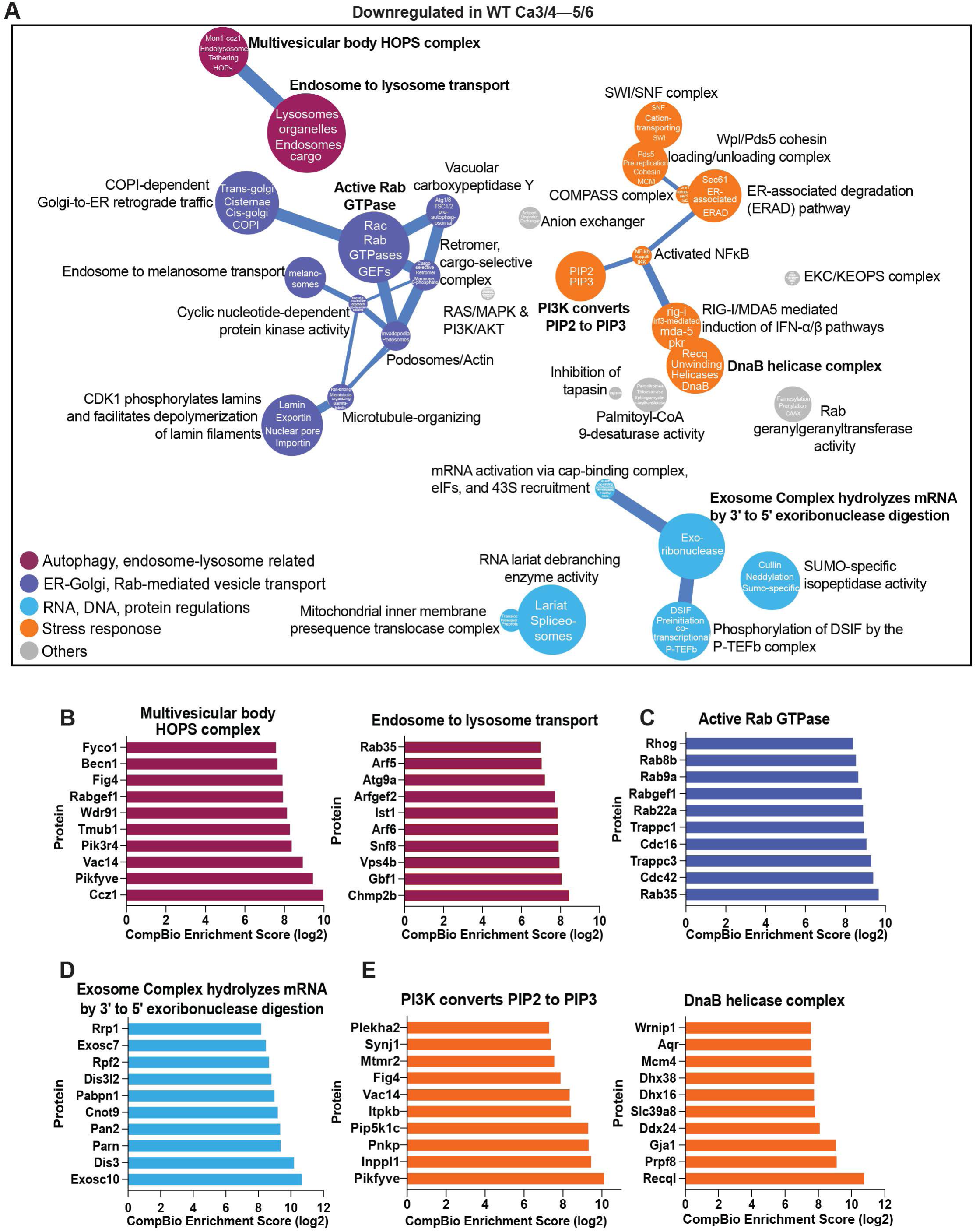
Frequent flexion-induced loading downregulates cellular stress response, including vesicle transport, autophagy, and RNA and DNA regulation in wildtype mice. (A) Biological themes associated with downregulated DEPs are highlighted. (B-E) CompBio analysis of DEPs and themes whose abundances were significantly downregulated under flexion-induced overloading.

### SDC4 is involved in regulating ciliary function, RNA and DNA stability in NP cells

To better understand the link between SDC4 and physiological disc loading, we compared genotype-dependent proteomic changes at spinal levels Ca6/7–8/9 (Fig. 6A). PCA revealed distinct clustering between WT and KO samples and showed 126 upregulated and 7 downregulated proteins in *Sdc4* KO (p ≤ 0.05, FC ≥1.5) (Fig. 6A’). Contextual analysis using CompBio identified four prominent upregulated thematic superclusters: 1) ciliary motility, 2) dynamin-regulated clathrin-coated cargo loading, 3) RNA regulation and DNA repair, and 4) other stress response themes (Fig. 6B). No downregulated themes were identified, as none met the statistical cutoff. Loss of SDC4 upregulated ciliary motility-related proteins, including CC2D2A, MKS1, IFT27, and IQCB1 (Fig. 6C). In parallel, themes related to dynamin- and Rab GTPase-mediated cargo transport and active lamellipodial cytoskeleton dynamics were upregulated (Fig. 6D-E). Proteins contributing to these themes included DNM3, RAB32, ARFGEF1, DENND1A, PIK3C2A, DIAPH2, DAAM1, ARFGEF1, and DOCK6 (Fig. 6D-E). This suggested that SDC4 is involved in regulating ciliary and membrane ruffling during the endocytic process by modulating cytoskeleton dynamics during the physiological loading of the spine.

**Fig. 6.**
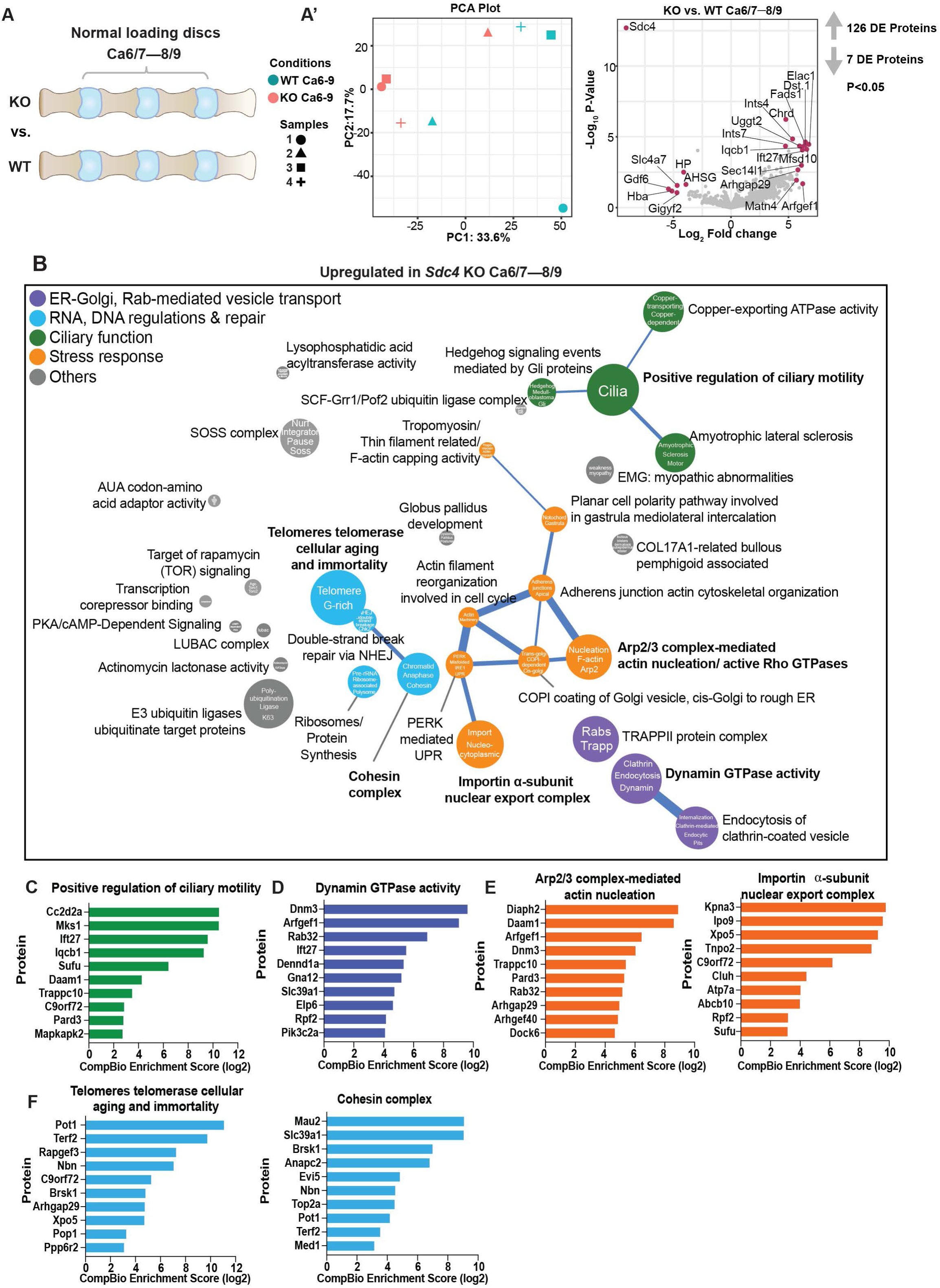
SDC4 regulates proteins associated with dynamin-mediated endocytic recycling processes and autophagy under normal loading condition. (A) Illustration comparing *Sdc4*-KO and WT NP under normal loading condition. (A’) The PCA plot shows distinct sample cluster separation, and the log_2_ fold-change volcano plot highlights DEPs that are up- and downregulated. (B) Enriched biological themes related to fast endocytic recycling, PERK-mediated UPR, and DNA regulation are highlighted. Downregulated themes did not meet the CompBio threshold cutoff. (C-F) Upregulated proteins and their associated biological themes.

Additionally, increased expression of KPNA3, IPO9, XPO5, TNPO2, and C9ORF72 within the importin alpha-subunit nuclear export complex theme suggested a heightened cellular adaptation to stress and enhanced RNA quality control. Concurrently, telomerase cellular aging and immortality and cohesin complex themes were associated with proteins such as POT1, TERF2, NBN, BRSK1, MAU2, ANAPC2, and TOP2A, suggesting potential regulation of DNA integrity and quality control mechanisms (Fig. 6F).

Next, to investigate the role of SDC4 in discs under increased spine flexion, we compared the proteome at Ca3/4–5/6 between genotypes (Fig. 7A). We identified 552 upregulated and 16 downregulated proteins (p ≤ 0.05, FC ≥ 1.5) (Fig. 7A’). While upregulated proteins showed a strong thematic clustering in CompBio, downregulated proteins did not. We uncovered four major thematic superclusters: 1) endosome-lysosomal autophagy-related pathways, 2) Rab-mediated intracellular vesicle trafficking, 3) stress response, and 4) RNA and DNA regulation (Fig. 7B). We observed enhanced activity of themes relating to Sec61 translocon complex/ERAD and CYLD-mediated deubiquitination of DDX58, with key proteins such as SEC63, UBE2J1, SEC61A2, RIG-I, IFIH1, DDX41, and G3BP1 (Fig. 7C). Moreover, proteins involved in COPI-dependent Golgi-ER retrograde trafficking and autophagosome assembly included VAMP4, STX3/8, SNX6, ZFYVE1, ATG9A, ATG3 (Fig. 7D-E). This increase in autophagosomes was supported by pronounced increase in LC3B puncta in *Sdc4*-KO discs (Suppl. Fig. 3). Lastly, RNA and DNA regulation-related proteins such as PLRG1, CPSF3, AQR, LIG4, APTX, NBN were enriched in themes such as spliceosomal tri-snRNP complex and non-homologous end-joining (NHEJ) pathway (Fig. 7F), suggesting a coordinated upregulation of nucleic acid repair and processing mechanisms under conditions of increased flexion. These findings suggested that SDC4 dampens autophagy by affecting the recycling of endosomes to lysosomes, potentially suppressing DNA damage repair and stress responses under high mechanical stress. Furthermore, using Assertion Engine (AE) functionality within CompBio we assessed the core function of SDC4 in NP cells regardless of their mechanical environment. We found shared concepts relating to Rab GTPases, clathrin-mediated vesicle trafficking, ER-golgi, actin and dynamin-mediated scission (Suppl. Fig. 4).

**Fig. 7.**
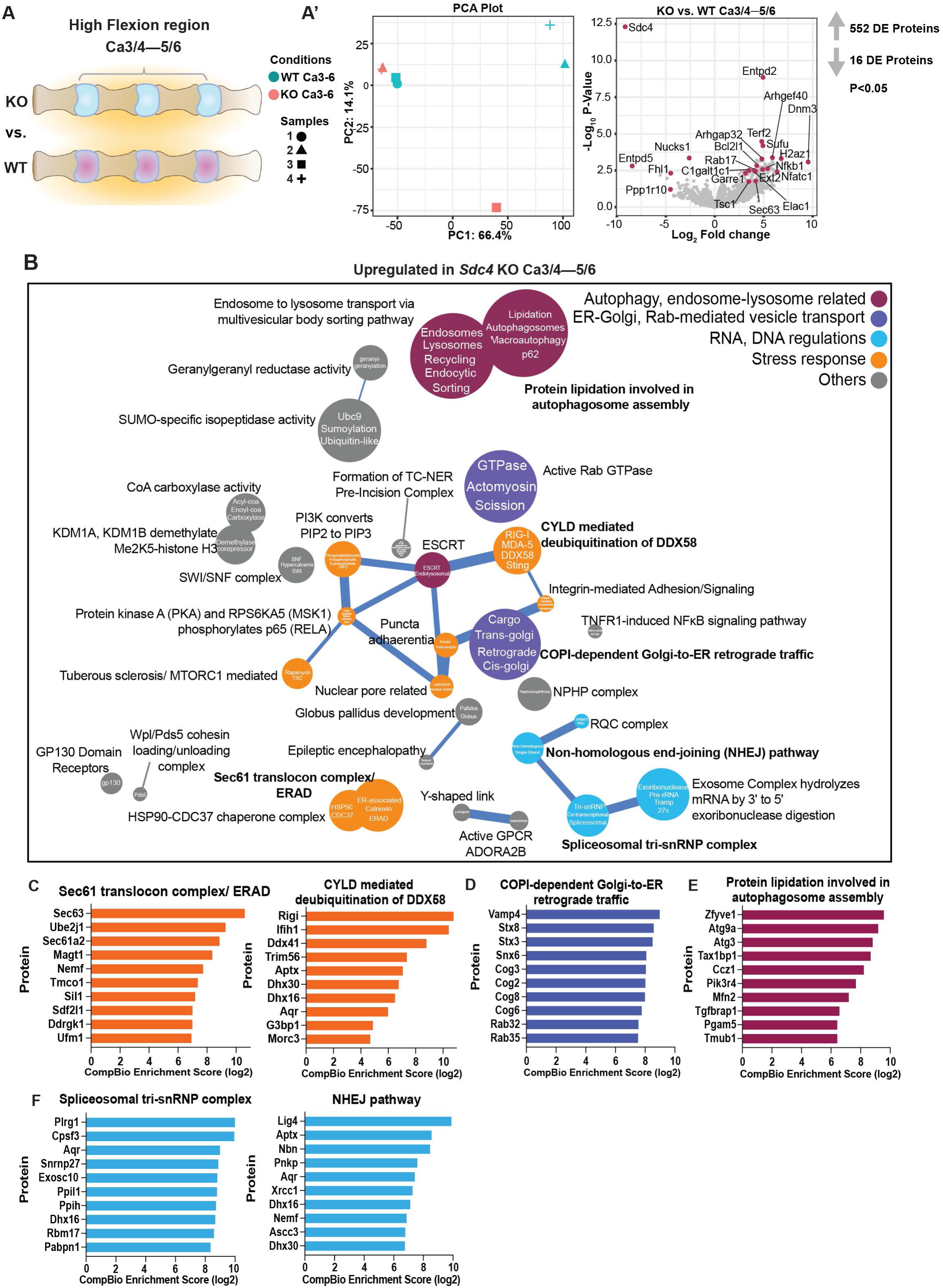
*Sdc4* KO NP upregulates endocytic recycling and autophagy related proteins to maintain cellular homeostasis under high flexion-induced stress. (A) Illustration comparing *Sdc4*-KO and WT NP under increase flexion-induced loading condition. (A’) The PCA plot shows sample cluster separation, and the log_2_ fold-change volcano plot highlights DEPs that are up- and downregulated. (B) Enriched biological themes related to stress response, Rab-mediated intracellular vesicle trafficking, and autophagy. Downregulated themes did not meet the CompBio threshold cutoff. (C-F) Upregulated proteins and their associated biological themes.

### Loss of SDC4 decreases disc cell apoptosis following acute mechanical stress

To investigate whether SDC4 loss mitigates loading-induced cell death, we used previously described tail looping model to apply asymmetric mechanical stress to the intervertebral discs Ca7-12 of WT and *Sdc4*-KO mice(23) (Fig. 8A). Histological analysis of Ca8-12 discs using Safranin O/Fast Green staining showed pronounced structural alterations in the overall disc architecture, as well as vertebral bone (VB) without overt differences between WT and KO mice (Fig. 8B), suggesting injurious nature of this loading regimen. To further investigate whether cells responded differently to this loading stress based on their SDC4 expression status, we measured cell apoptosis. TUNEL staining showed that SDC4 deletion significantly decreased the percentage of apoptotic cells in both the AF and NP regions compared to WT controls (Fig. 8C-E). Next, we examined regional differences within the AF compartment, the side that experienced either compressive or tensive loading. On the compressed side, the percentage of TUNEL-positive cells was comparable between the genotypes (Fig. 8F). Interestingly, however, in the AF region with increased tension, *Sdc4*-KO mice exhibited a significant reduction in TUNEL-positive cells compared to WT (Fig. 8G). Our findings suggested that *Sdc4* deletion protects the discs against mechanical loading-induced apoptosis, suggesting a role for SDC4 in stress adaptation.

**Fig. 8.**
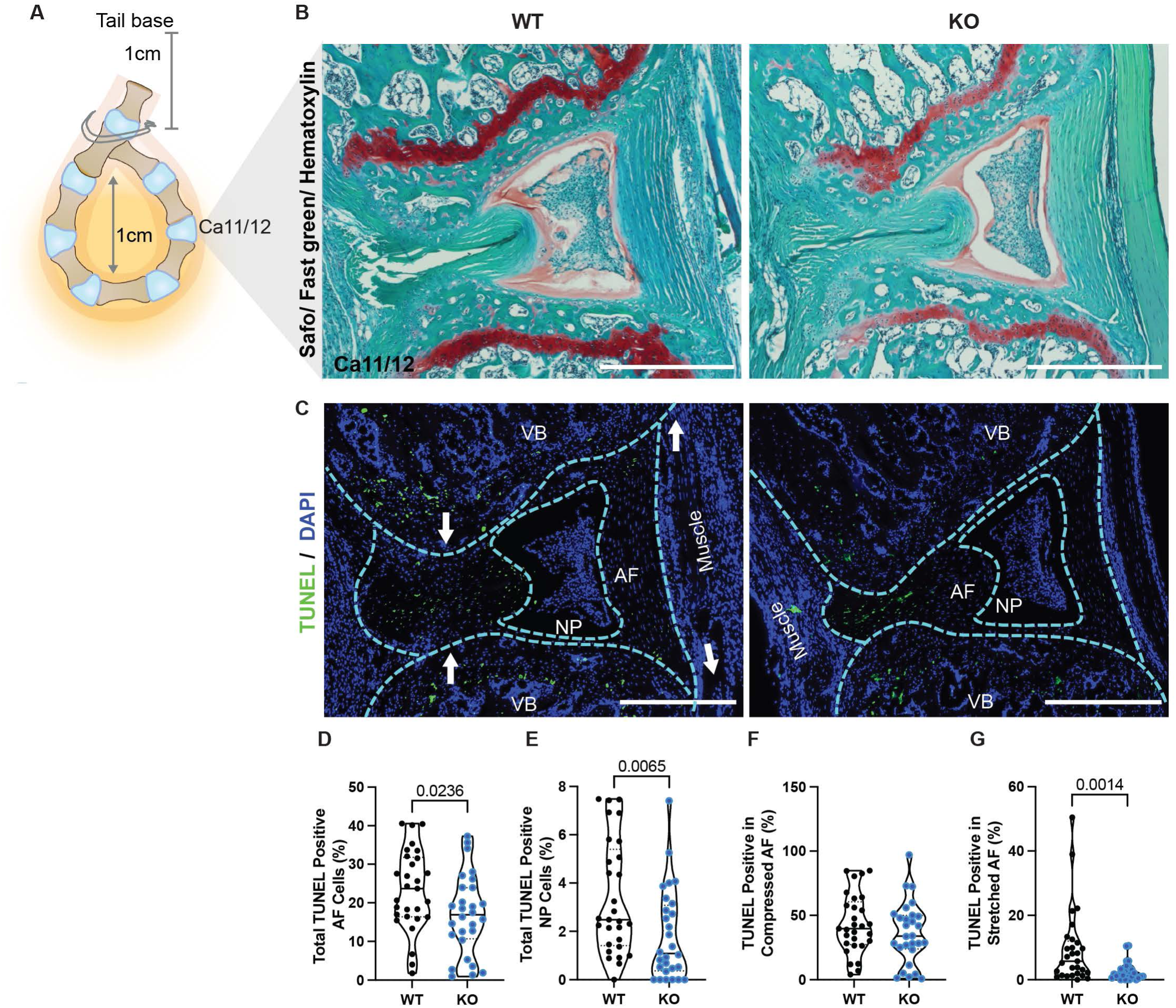
*Sdc4-*KO discs mitigate cell death under tail looping-induced mechanical loading. (A) Illustration showing tail looping 1cm from the tail base with a 1cm loop diameter. (B) Safranin-O/Fast Green/Hematoxylin staining of WT and *Sdc4-*KO looped discs. (C) TUNEL staining of WT and *Sdc4-*KO looped discs. (D-E) Quantification of total TUNEL-positive cells in the AF and NP. (F-G) Quantification of TUNEL-positive AF cells under compressive and tensive stress. Ca8/9–12/13 discs were examined. 9 WT mice (5M, 4F), 29 discs total; 7 KO mice (5M, 2F), 29 discs. Scale bar, 250μm. Violin plots show score distribution with quartile range and median. Significance was determined using an unpaired Welch’s t-test or Mann-Whitney test, P < 0.05.

### Loss of SDC4 compromises vertebral bone health in the caudal spine

Although SDC4 loss mitigated mechanical loading-induced disc degeneration, its effects on vertebral bone mineral density and health merits further consideration. Therefore, we examined bone health parameters of Ca3–6 vertebrae from WT and *Sdc4*-KO mice. We observed pronounced reductions in trabecular bone structural parameters such as bone volume (BV/TV), trabecular number, spacing, thickness, and bone mineral density in the KO group; in cortical bone, KO mice also showed decrease in mineral density (Suppl. Fig. 5A-C).

## DISCUSSION

The mechanical environment of musculoskeletal tissues is a critical determinant of tissue homeostasis, regeneration, and pathology(24–27). Recent reports suggest that increased flexion near the base of the mouse caudal spine accelerates early disc degeneration(5, 28). Given our previous findings that SDC4 controls matrix homeostasis by regulating ECM secretion and collagen cross-linking(7), we investigated whether SDC4 loss mitigates early degenerative processes by limiting ECM deposition and remodeling. Our findings provide compelling evidence that SDC4 promotes the early-onset disc degeneration when subjected to increased flexion by affecting NP cell phenotype as evidenced by a transition from TAGLN-positive to COL X-positive hypertrophic NP cells. The accumulation of denatured collagen chains and FN further underscored the dysregulated functionality of these cells(29–31). Additionally, histological grading and FTIR analyses highlighted a stark difference in morphology and collagen cross-linking between *Sdc4*-KO and wild-type discs, underscoring the pivotal role of SDC4 in mediating fibrotic remodeling under altered mechanical stress. These observations aligned with previous findings that demonstrated the interaction of SDC4 with lysyl oxidase (LOX), FN, and transglutaminase 2 (TGM2) to promote collagen fiber formation and facilitate collagen cross-linking and fibrosis(8, 32, 33).

At the cellular level, mechanical cues can trigger cytoskeletal rearrangements and activate signaling pathways, thereby influencing downstream gene transcription and cellular adaptation to stressors(34, 35). Herein, NP tissues subjected to altered loading showed increased levels of chondrogenic markers GDF5 and COL10A1, which are pivotal for cartilage matrix homeostasis and cellular phenotype regulation, suggesting that NP cells have acquired a more chondrocytic phenotype(36, 37). Additionally, elevated expression of NEB, TPM2, TTN, ACTN2 suggests enhanced cytoskeletal reorganization and contractility. Several studies demonstrated the importance of cellular contractility and tissue contraction in cellular adaptation to mechanical cues(38, 39). Moreover, upregulation of PLOD2, CCN2, TIMP3, EFEMP2/fibulin-4 suggested an active ECM remodeling in the disc by promoting collagen stability and cross-linking(17, 18, 40).

Our analysis also uncovered that altered mechanical loading causes a significant downregulation of proteins involved in regulating vesicular transport, ERAD, and mRNA quality. For example, the reduction of vesicle transport regulators—CCZ1, VAC14, and PIKFYVE— suggested a compromised ability of NP cells to sustain proteostasis. Notably, CCZ1 and its partner MON1 form a guanosine exchange complex that activates RAB5 and RAB7 GTPases, which are critical for autophagosome-lysosome fusion, intracellular trafficking, and overall organismal development(41, 42). In parallel, the kinase PIKFYVE and the phosphatase FIG4 assemble with the scaffolding protein VAC14 to control phosphatidylinositol 3,5-bisphosphate (PI(3,5)P2) metabolism(43–45). This lipid not only serves as a key signaling molecule but also acts as a precursor for phosphatidylinositol 5-phosphate (PI5P), a crucial messenger in stress response, vesicular trafficking, and cell survival(46). Beyond proteostasis, excessive mechanical stress also appears to compromise RNA integrity, as evidenced by the reduction in RNA exosome complex-related proteins EXOSC10, DIS3, and PARN. Given the central role of exosomes in RNA processing, decay, and surveillance, its impaired function may disrupt transcriptomic stability, further exacerbating cellular dysfunction(47, 48). Accordingly, the impairment of essential stress response mechanisms in NP cells likely accelerates cellular deterioration while at the same time engaging in adaptive response aimed at reinforcing the ECM and restoring tissue mechanical integrity. These findings underscore the complex interplay between anabolic and catabolic processes in the pathogenesis of disc degeneration and are consistent with prior studies linking excessive mechanical stress to dysregulated ECM turnover and impaired cellular stress responses(49–51).

Notably, we observed that SDC4 was involved in cytoskeleton dynamics, vesicular trafficking, and RNA and DNA regulation in NP cells. The upregulation of ciliary motility-related proteins, including CC2D2A, MKS1, and IFT27 in *Sdc4-*KO NP points to a possible role for SDC4 in regulating cilia function, critical for cellular signaling and mechanosensing in the disc(52–54). Cells constantly remodel their cytoskeleton to respond to external cues, and SDC4 may play a role in fine-tuning this adaptability. Supporting this, several key regulators of cytoskeletal reorganization such as DNM3, RAB32, PIK3C2A, DAAM1, and DOCK6 were upregulated. These proteins orchestrate membrane ruffling through RHOG-, CDC42-, and RAC1-mediated signaling while also facilitating endocytic trafficking via DNM3(55–60). This coordinated response suggested that in NP cells, SDC4 may fine-tune cytoskeletal dynamics to maintain cellular stability(61). While our data suggested that *Sdc4* deletion increases endocytic process, Bass *et al.* reported that disruption of SDC4, caveolin, or RhoG in mice delays closure of dermal wounds due to a migration defect of the fibroblasts(62). We speculate that these differences in response mechanism arise due to the role that syndecans play in different tissue niches and specific cell types. Furthermore, increased RNA-regulating proteins, including KPNA3, XPO5, and C9ORF72, are integral to nucleocytoplasmic transport, RNA toxicity regulation, and quality control(63). This may be part of an adaptive response to counteract the potential disruptions in cellular homeostasis, underscoring the efforts to maintain protein synthesis and RNA stability under stress.

SDC4 similarly impacted stress response, intracellular vesicle trafficking, and RNA and DNA regulation in discs exposed to increased flexion. Notably, increased expression of proteins involved in ERAD (SEC63, UBE2J1, SEC61A2) and key regulators of the autophagosome assembly (ZFYVE1, ATG9A, and ATG3) suggested enhanced protein and organelle quality control in *Sdc4* KO discs(64–67). Furthermore, increased levels of PLRG1, CPSF3, and LIG4, suggested enhanced nucleic acid processing and repair, likely to mitigate the DNA damage induced by increased flexion-induced stress(68–70). While there are no current reports on SDC4 regulating RNA and DNA repair, we speculate that SDC4 loss activates autophagy(71) and supports autophagy-mediated protection against genetic instability, thus indirectly affecting DNA and RNA quality control(72). Furthermore, decreased incidence of cell death in NP and AF compartments of tail-looped discs suggested that SDC4 loss partially counteracted mechanical overloading-induced cell apoptosis. However, a key limitation of this model is the static nature of the applied mechanical load, which may elicit an aggressive cellular response compared to the dynamic flexion-induced stress paradigm.

It is important to comment here on the low vertebral bone mass phenotype we noted in KO mice. While the causal relationship between vertebral osteopenia and disc degeneration remains uncertain in clinical studies, it is clear that altered loading dynamics can impact the overall health of the spinal motion segment(73, 74). For example, a recent finite element study found that von Mises stress on the NP is more concentrated in osteoporotic spines compared to normal ones, potentially accelerating disc degeneration(75). Therefore, the protective effect and structural changes we observed in the discs of *Sdc4* KO mice are unlikely to be secondary effects of the osteopenic vertebral bone.

In summary, our work shows that SDC4 fine-tunes mechanical stress response in disc cells by reorganizing their cytoskeleton through membrane ruffling, a dynamic process driven by actin polymerization that enhances extracellular sensing and uptake of signaling molecules. It is possible that these changes facilitate the internalization of ECM components and stress-responsive factors, priming cells for downstream vesicular trafficking and degradation processes. In mechanically stressed NP cells, dysregulation of these vesicular transport regulators impairs proteostasis, compromising autophagosome-lysosome fusion. However, in *Sdc4* KO discs, enhanced autophagy-related protein levels may indicate the presence of a compensatory mechanism that maintains protein and organelle quality control, indirectly promoting RNA and DNA quality control to ensure transcriptomic stability. Thus, SDC4 is an important component of mechanosensing machinery that elicits a compensatory and remodeling response to altered mechanical loading in the intervertebral disc.

## MATERIALS AND METHOD

### Mice

All animal care procedures, housing, breeding, and the collection of animal tissues were performed in accordance with a protocol approved by the Institutional Animal Care and Use Committee (IACUC) of Thomas Jefferson University. The *Sdc4* knockout mice on C57BL/6 background have been described earlier(76). These mice harbor deletions of exon 2 to part of exon 5 that encodes the N-terminal coding region. Both male and female mice were used in these studies.

### Histological Analysis

Caudal spines from 6-month-old mice were dissected and immediately fixed in 4% PFA in PBS at 4°C for 48h, decalcified in 20% EDTA at 4°C for 2-3 weeks, and then embedded in paraffin. 7µm mid-coronal sections were cut from Ca3-6 and used for histological staining. Sectioned tissues were deparaffinized, followed by graded ethanol rehydration preceded all staining protocols. Following Safranin-O/Fast Green/Hematoxylin staining, images were acquired on a light microscope (Axio Imager 2; Carl Zeiss Microscopy) using 5x/0.15 N-Achroplan (Carl Zeiss) objective and Zen2™ software (Carl Zeiss). The health of disc compartments was assessed by at least four blinded graders using Modified Thompson Grading. Picrosirius red staining (Polysciences, 24901) was performed to assess collagen fibril thickness, and images were acquired using 4x /0.25 Pol /WD 7.0 (Nikon) objective on a polarizing light microscope (Eclipse LV100 POL; Nikon). NIS Elements Viewer software (Nikon) was used to set color threshold for the level of green (thin), yellow (intermediate), and red (thick) fibers. Color threshold levels remained constant for all samples. For micro-dissection observation of the caudal spines, NP tissue was observed under light microscope (Carl Zeiss), absent or presence of fibrosis was noted based on the texture of the tissue. Gel-like NP tissue was scored as absent, while fibrotic NP tissue was scored present.

### Imaging Fourier-Transform Infrared Spectroscopy

7µm deparaffinized sections of decalcified caudal disc (Ca3-6) were used to acquire infrared (IR) spectral imaging data using methods previously described(7).

### Immunohistochemistry and digital image analysis

Mid-coronal sections (Ca3-6) were deparaffinized in histoclear and rehydrated in ethanol solutions (100-95%), water, and PBS. Antigen retrieval using citrate-buffer at pH 6 (Vector Laboratories) was performed on samples stained with COL I (1:100; Abcam, ab34710), COL II (1:400; Fitzgerald, 70R-CR008), CS (1:300; Abcam, ab11570), TAGLN (1:100; Abcam, ab14106), FN (1:100; NBP1-91258), and LC3B (1:100; NB100-2220). Enzymatic antigen retrieval was used for COLX (1:500; Abcam, ab58632; 1:1000 Proteinase K), and ACAN (1:50; Millipore Sigma, AB1031; 1:200 chondroitinase ABC (20U/mL)). MOM kit (Vector Laboratories; BMK-2202) was used per manufacturer instruction. After antigen retrieval, samples were blocked with 5-10% normal goat or donkey serum (Jackson ImmunoResearch) in PBS-Triton (0.4% Triton X-100 in 1x PBS) for 1h at room temperature. Primary antibodies were applied and incubated overnight at 4°C. Samples were then generously washed three times with PBS and incubated with Alexa Fluor-594-conjugated secondary antibody (1:700, Jackson ImmunoResearch) for 1h at room temperature, shielded from light. Samples were washed three times with PBS and mounted with ProLong™ diamond antifade mountant with DAPI (Invitrogen, P36971). Images were acquired on Axio Imager 2 (Carl Zeiss Microscopy) using either 5×/0.15 N-Achroplan (Carl Zeiss),10×/0.3 EC Plan-Neofluar (Carl Zeiss) objective or 20×/0.5 EC Plan-Neofluar (Carl Zeiss), X-Cite 120Q Excitation Light Source (Excelitas Technologies), AxioCam MR R3 camera and Zen 2TM software (Carl Zeiss Microscopy). Per staining experiment, images were taken at a set exposure time for all samples. LC3B staining was captured on a Zeiss LSM 800 Axio Inverted confocal microscope (Plan-Apochromat 63x/1.40 oil). All quantifications were done on ImageJ 1.52i (NIH). Images were thresholded to create binary images, and NP and AF compartments were manually contoured using the Freehand Tool and ROI were analyzed using Area Fraction measurement tool. LC3B puncta were counted using the Analyze Particles tool and each cell is manually counted by DAPI marker; puncta per cell were then calculated.

### Denatured collagen hybridizing peptide (CHP) assay

Mid-coronal Ca3-6 disc sections (7μm) from WT and KO mice, (6 animals, 3 discs/animal) were deparaffinized in histoclear and rehydrated in ethanol solutions (100-95%) and water. F-CHP (3Helix, FLU60) was applied according to the manufacturer’s protocol. Sections were washed and mounted, and images were acquired as described above.

### TUNEL assay

TUNEL-staining was performed using the Click-iT™ Plus TUNEL Assay Kits for In Situ Apoptosis Detection Kit (Thermo Fisher scientific, C10618). Briefly, sections were de-paraffinized and permeabilized using Proteinase K (20 μg/mL) for 15 min at room temperature and a TUNEL assay was carried out per the manufacturer’s protocol. Sections were washed and mounted with ProLong™ Diamond antifade mountant with DAPI. TUNEL-positive analysis was done as described for LC3B.

### Protein extraction and LC-MS/MS analysis

Each sample contains six NP tissues. Per mechanically overloaded sample condition, three caudal discs (Ca3/4, Ca4/5, Ca5/6) were pooled from two animals; and per normal loading sample condition, three contiguous caudal discs (Ca6/7, Ca7/8, Ca8/9) from the same two animals were pooled. Samples were scooped into 200µL of chaotropic buffer (pH 5.8) containing 50mM Sodium acetate (NaAc), 4M Guanidine hydrochloride (GdnHCl), 100mM 6-aminocaproic acid, 5mM benzamidine and 5mM N-ethylmaleimide, sonicated on ice (two cycles of 5 s bursts with intensity at 40% and 5 s breaks). Additional 200µL of chaotropic buffer was added to each sample and proteins were extracted at 4°C on rotator for 48h. Soluble proteins were collected after centrifugation at 13,200xg, 4°C for 30 min, and reduced with 4mM DTT at 56°C for 1h, alkylated with 50mM iodoacetamide at room temperature in the dark for 1h, and reaction terminated by ethanol precipitation at 4°C overnight. Proteins were recovered by centrifugation and pellets dried under vacuum and solubilized in 4M urea, and protein content quantified using the BCA assay(14). Proteins were submitted to Wistar Proteomics and Metabolomics Facility for LC-MS/MS analysis. 15µg of each sample were run into 10% NuPAGE Bis-Tris gels using given concentrations. Each sample/lane was reduced with TCEP, alkylated with iodoacetamide, and digested in-gel with trypsin. The tryptic digest of each sample was then analyzed using a 3h LC-MS/MS DIA (OTOT) run on the Thermo Orbitrap Astral mass spectrometer. LFQ-Analyst was used to analyze differentially expressed proteins (https://analyst-suite.monash-proteomics.cloud.edu.au/apps/lfq-analyst/). Matrisome analysis was done using Matrisome AnalyzeR(15). Contextual analysis was done using CompBio.

### Proteomic data analyses using CompBio tool

Differential protein lists from NP tissues of *Sdc4* KO mice (1.5-fold change, P-value < 0.05) were analyzed using CompBio (V2.0, PercayAI Inc., www.percayai.com/compbio)(7). CompBio is an AI-driven platform designed to interpret multi-omics data through a systematic analysis of biological entities, such as genes, proteins, miRNAs, or metabolites, based on literature from PubMed. The platform processes both abstracts and full-text articles using natural language processing and conditional probability analysis to construct an extensive, up-to-date knowledge base. CompBio evaluates all input entities for statistically enriched biological “concepts” and clusters these into broader “themes” based on their co-occurrence patterns within PubMed articles. The relationships between themes are displayed on an interactive knowledge map, where themes are represented as spheres. The connections between themes (edges) reflect shared biological entities, with thicker edges indicating a greater number of shared genes.

Themes are ranked by their enrichment scores, with the sphere size representing the relative enrichment based on the absolute enrichment score. Each theme is assigned a normalized enrichment score (NES) and P-value, determined by comparing the score distribution against randomized datasets. Significant themes (NES > 1.3 and P-value < 0.1) are further reviewed for biological interpretation. Unlike conventional methods, CompBio does not require stringent fold-change or P-value cutoffs to filter out noise, as its holistic analysis approach accounts for the entire experimental context. The platform annotates each theme based on concept enrichments and their relationships with adjacent themes, providing a comprehensive, experiment-aware interpretation. Additionally, since related biological processes may produce multiple interconnected themes, each theme can be labeled with up to three biological descriptors to fully represent its underlying components. For global similarity calculations, we used Assertion Engine (AE V 2.4.3) to analyze two datasets at the concept level as well as compares the interrelationships of the conserved concepts. If the same enriched concept is present in both datasets, the AE determines how similar or different that concept’s relationships are with the other concepts in the respective datasets. A score of 0.0 represents no contextual biological similarity whereas a score of 1.0 represents complete similarity.

### Tail looping

Tail looping was previously described to induce disc degeneration(23). 6-month-old WT and KO animals were anesthetized with 3-4% isoflurane. A mark was placed 1cm from the tail base to ensure that the Ca3/4–5/6 region was avoided. Using a simple non-surgical procedure, the remaining caudal spine was then looped at a fixed position with a 0.8-mm stainless steel wire (Thermo Scientific, AA43301CC). To minimize excessive variability in stress on the discs, each tail was looped to maintain a consistent 1cm diameter measured from cranial to caudal position. Animals were then collected for histology at 14 days after looping.

### Micro-Computed Tomography (µCT) Analysis

Caudal spines from *Sdc4*-KO and WT mice were dissected and fixed in 4% paraformaldehyde (PFA) in PBS at 4°C for 48h before performing micro-CT (µCT) scans (Bruker, Skyscan 1275). To evaluate bone morphometry, the spines were rinsed, rehydrated in PBS, and scanned at a resolution of 15µm^3^ per voxel (50kV, 200µA, 85ms exposure time, 0.2° rotation step) using a 1mm aluminum filter. Image reconstruction was carried out with the Skyscan NRecon software package. Trabecular bone parameters were quantified using Skyscan CT analysis (CTAn) software by contouring the region of interest (ROI) within the 3D-reconstructed trabecular tissue. The following parameters were measured: bone volume fraction (BV/TV), trabecular number (Tb.N), trabecular thickness (Tb.Th), and trabecular separation (Tb.Sp).

Cortical bone analysis was performed in two dimensions, with measurements taken for bone volume (BV), cross-sectional thickness (Cs.Th), mean cross-sectional bone area (B.Ar), and mean cross-sectional tissue area (T.Ar). Mineral density was determined using a standard calibration curve generated with a pair of mineral density phantoms (0.25g/cm^3^ and 0.75g/cm^3^ calcium hydroxyapatite (CaHA)).

### Statistical analysis

Statistical analysis was performed using Prism 10 (GraphPad, La Jolla, CA, USA) with data presented as violin plots showing all data points with median and interquartile range and maximum and minimum values. Differences between distributions of two samples were checked for normality using Shapiro–Wilk tests and further analyzed using an unpaired Welch’s t-test for normally distributed data and the Mann–Whitney U test for non-normally distributed data. Two-way ANOVA with Tukey’s multiple comparison was used to determine the effects on interaction between genes and disc levels. Since interactions between genetic, biological, and biomechanical factors at individual spinal levels have been shown to produce different phenotypic outcomes, each intervertebral disc or vertebra is considered an independent sample a methodology followed by many groups(77, 78). Analyses of Modified Thompson Grading data distributions and fiber thickness distributions were performed using a chi-square test at a 0.05 level of significance.

## ACKNOWLEDGMENTS

We would like to thank Dr. Hsin Yao Tang and Oreoluwa Solanke at the Wistar Institute Proteomics & Metabolomics Facility for providing proteomic service. Illustrations were created with BioRender.com.

## AUTHOR CONTRIBUTIONS

Kimheak Sao: Conceptualization; methodology; investigation; data curation; formal analysis; validation; visualization; writing– original draft; writing – review and editing. Makarand V Risbud: Conceptualization; funding acquisition; resources; validation; supervision; writing– review and editing.

## DATA AVAILABILITY STATEMENT

The mass spectrometry proteomics RAW data have been deposited to the ProteomeXchange Consortium via the PRIDE partner repository with the dataset identifier PXD061689.

## ETHICS STATEMENT

All animal experiments were performed under IACUC protocols approved by Thomas Jefferson University.

## Author contributions

K.S. Conceptualization; methodology; investigation; data curation; formal analysis; validation; visualization; writing– original draft; writing – review and editing. M.V.R: Conceptualization; validation; funding acquisition; resources; supervision; writing– review and editing.

---

**Suppl. Fig.1.**
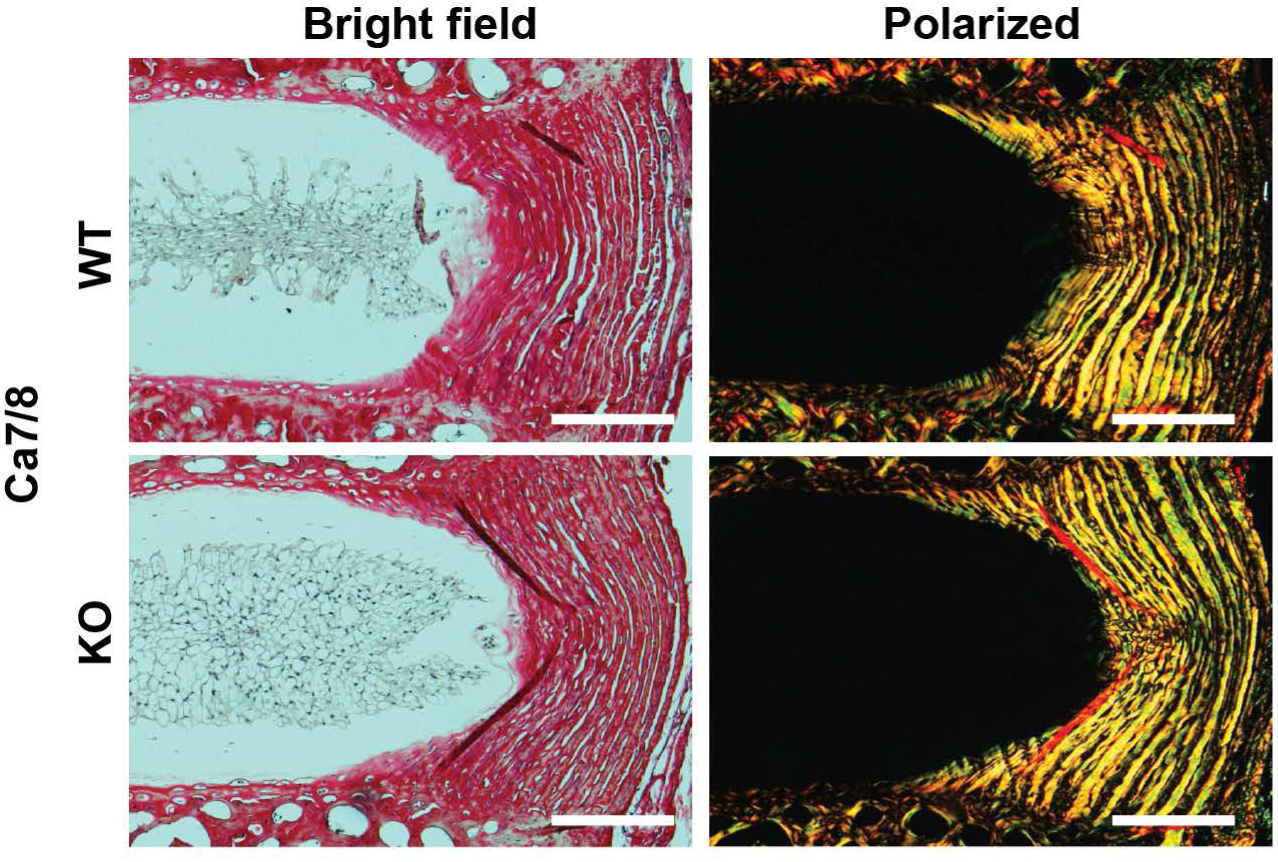

**Suppl. Fig.2.**
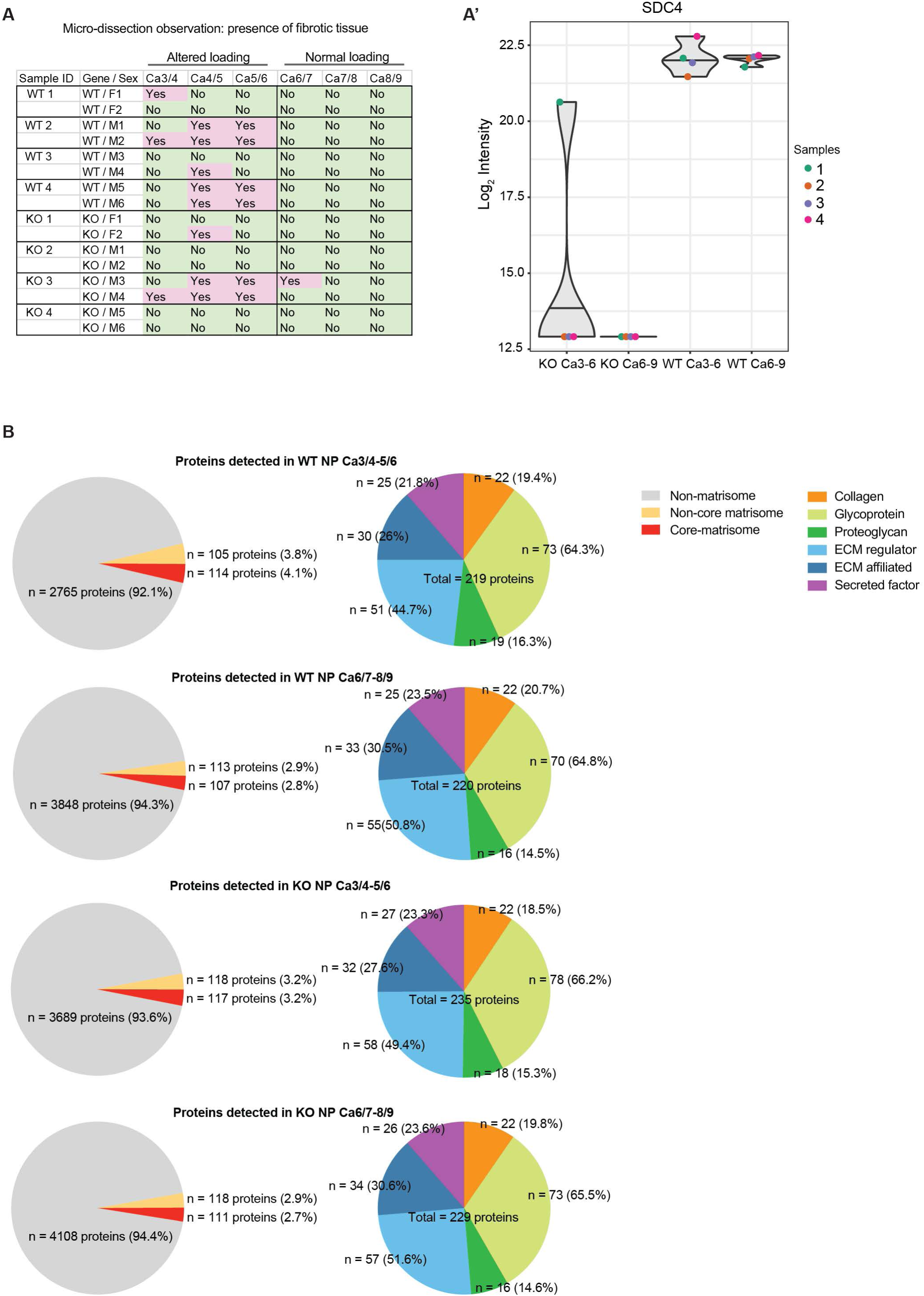

**Suppl. Fig.3.**
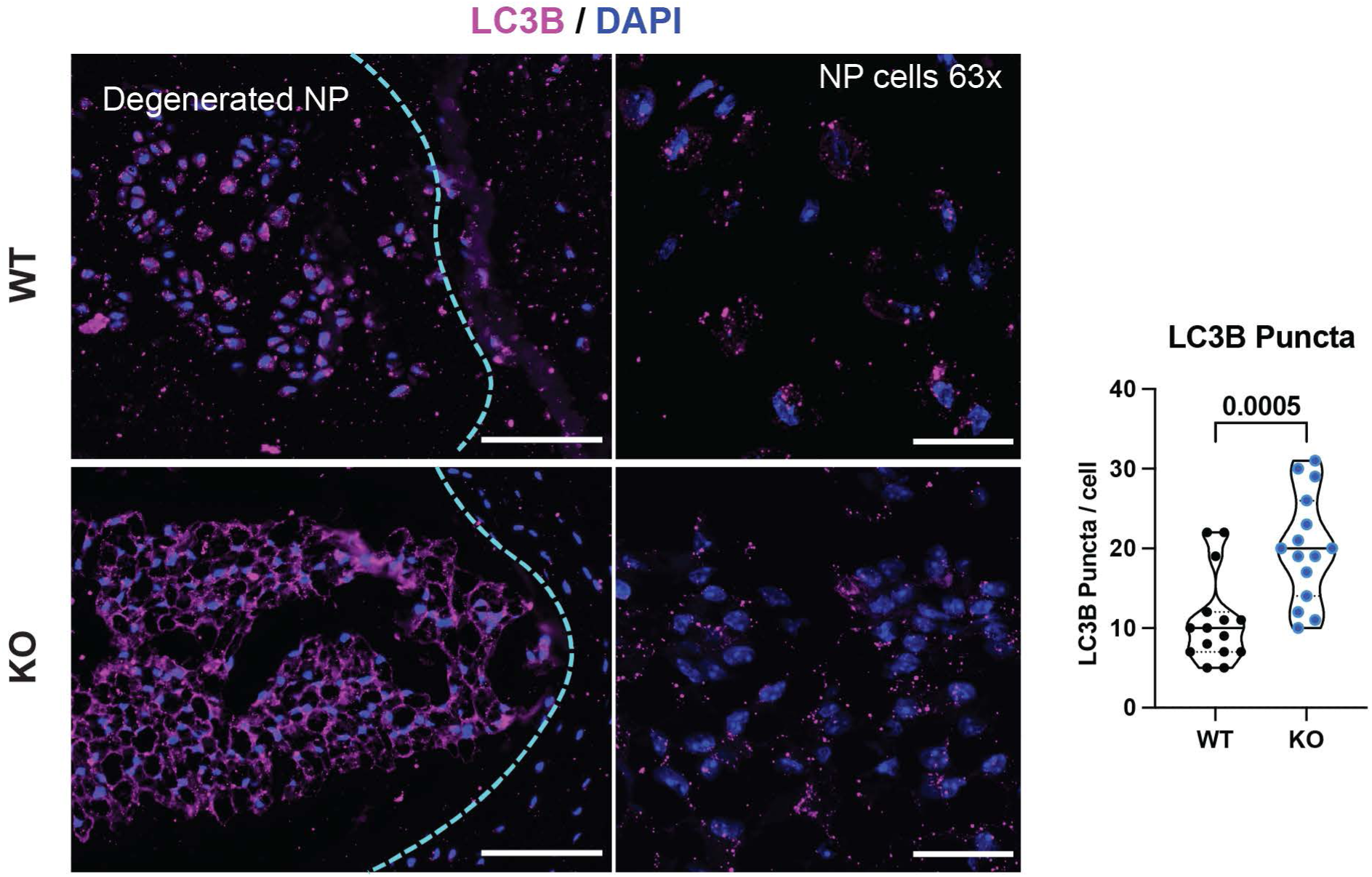

**Suppl. Fig.4.**
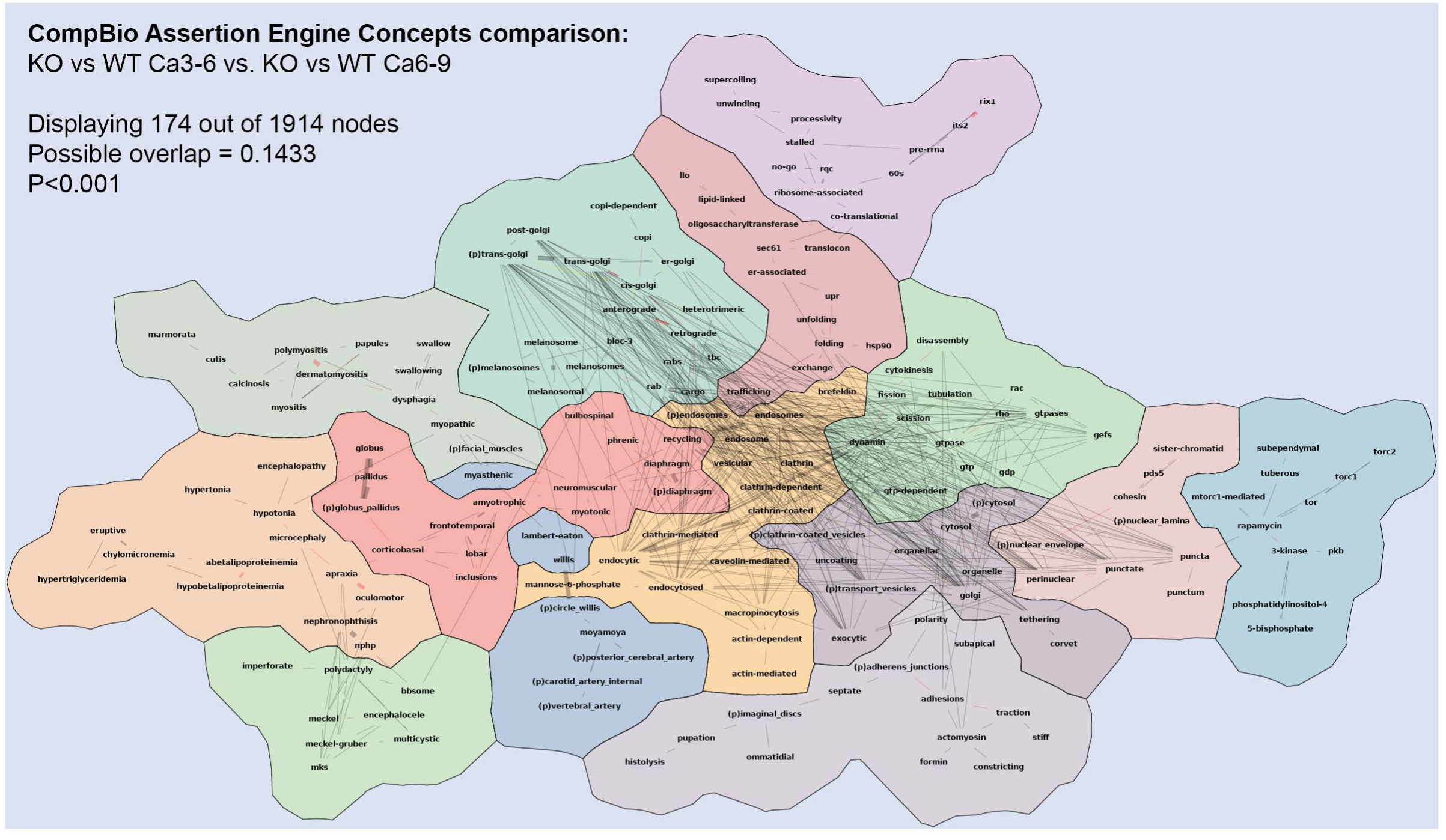

**Suppl. Fig.5.**
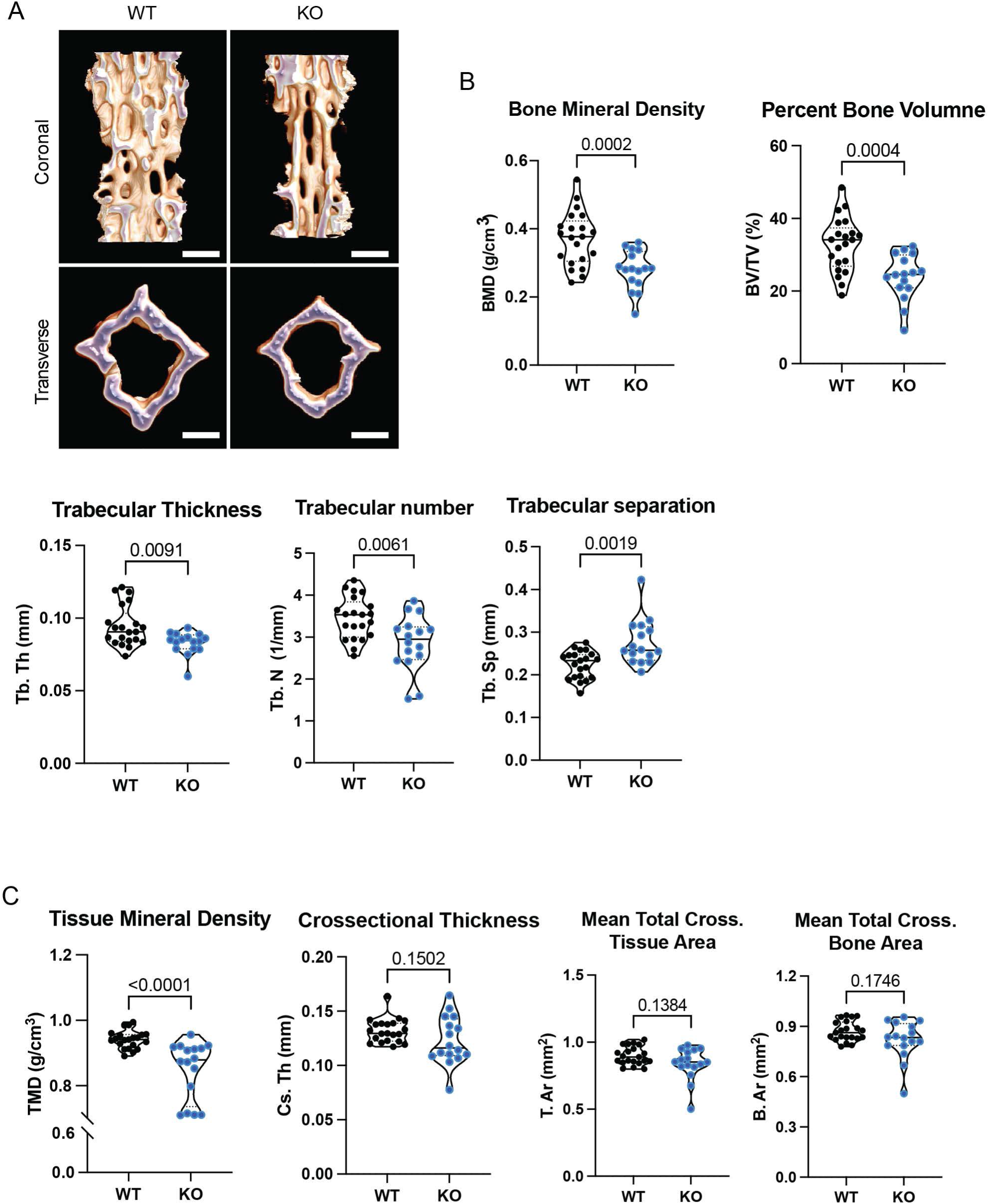

