## Supplementary material for "SDC4 drives fibrotic remodeling of the intervertebral disc under altered spinal loading": Suppl. Figure Legends

**SUPPLEMENTAL FIGURE LEGENDS**

**Supplemental Fig. 1. Picrosirius red staining of mouse caudal discs that experience normal loading.** Representative brightfield and corresponding polarized images of Ca7/8 disc from *Sdc4*-KO and WT showing lack of collagen deposition in the NP compartment.

**Supplemental Fig. 2. Description of samples used for proteomics and identified protein categories.** (A) Description of NP tissue samples used for proteomic analysis (A') SDC4 protein signal determined by LC-MS/MS in Ca3/4–5/6 and corresponding Ca6/7–8/9 discs from each sample. KO Ca3/4–5/6 sample 1 was determined to be an outlier; and omitted from the downstream analysis. (B) Detail categorization and number of proteins in non-core matrisome and core matrisome from WT and KO discs.

**Supplemental Fig. 3 LC3B staining in the NP compartment.** (A) Representative images showing localization of autophagosome marker LC3B in WT and KO NP tissues. Quantification of LC3B positive autophagosome puncta per cell. 5 WT mice (5M), 15 discs; 7 KO mice (6M, 1F), 15 discs. Violin plots show score distribution with median and quartile range. Significance was determined using an unpaired Mann-Whitney test, $P < 0.05$.

**Supplemental Fig. 4 CompBio Assertion Engine analysis of proteome showing shared SDC4-dependent concepts between Ca3-6 and Ca6-9 discs.** (A) Territorial map showing the preserved themes/concepts that are SDC4-dependent irrespective of the mechanical loading environment of the spine.

**Supplemental Fig. 5 *Sdc4*-KO caudal vertebrae experience early osteopenia.** (A) Representative 3D rendered trabecular and cortical tissue shows thinning of the vertebral bone. Scale bars: 0.5mm. (B) Bone mineral density (BMD) ($g/cm^3$), Percent bone volume/ tissue volume (BV/TV) (%), trabecular thickness (Tb.Th) (mm), trabecular number (Tb.N) (1/mm), trabecular separation (Tb.Sp) (mm) in trabecular tissue were analyzed. (C) Measurements of cortical bone parameter showing tissue mineral density (TMD) ($g/cm^3$), cross-sectional thickness (Cs.Th) (mm), mean total cross-sectional thickness bone area (B.Ar) ($mm^2$), mean total cross-sectional tissue area (T.Ar) ($mm^2$). At vertebrae Ca4-5, 11 WT mice (8M, 3F), 21 vertebrae and 10 KO mice (9M, 1F), 16 vertebrae were analyzed. Violin plots show score distribution with median and quartile range. Significance was determined using an unpaired Welch's t-test or Mann-Whitney test, $P < 0.05$.

**Table 1. CompBio concepts, themes, and their associated proteins.** A complete list of Down- and Up-regulated themes and concepts mapped by CompBio analysis with Normalized Enrichment Score > 1.2 & p-value < 0.1.

**Table 2. Matrisome analysis of total proteins extracted.** A complete list of core-matrisome and non-core matrisome annotation of total proteins extracted.
